# Quantifying the Brain Predictivity of Artificial Neural Networks with Nonlinear Response Mapping

**DOI:** 10.1101/2020.09.27.315747

**Authors:** Aditi Anand, Sanchari Sen, Kaushik Roy

**Affiliations:** West Lafayette Jr/Sr High School, West Lafayette, IN, USA; IBM T.J. Watson Research Center; Center for Brain-inspired Computing, School of Electrical and Computer Engineering, Purdue University, West Lafayette, IN, USA

**Keywords:** Artificial neural networks, brain-inspired computing, brain similarity, neural recordings, neural response predictivity

## Abstract

Quantifying the similarity between artificial neural networks (ANNs) and their biological counterparts is an important step towards building more brain-like artificial intelligence systems. Recent efforts in this direction use *neural predictivity*, or the ability to predict the responses of a biological brain given the information in an ANN (such as its internal activations), when both are presented with the same stimulus. We propose a new approach to quantifying neural predictivity by explicitly mapping the activations of an ANN to brain responses with a nonlinear function, and measuring the error between the predicted and actual brain responses. Further, we propose to use a neural network to approximate this mapping function by training it on a set of neural recordings. The proposed method was implemented within the Tensorflow framework and evaluated on a suite of 8 state-of-the-art image recognition ANNs. Our experiments suggest that the use of a non-linear mapping function leads to higher neural predictivity. Our findings also reaffirm the observation that the latest advances in classification performance of image recognition ANNs are not matched by improvements in their neural predictivity. Finally, we examine the impact of pruning, a widely used ANN optimization, on neural predictivity, and demonstrate that network sparsity leads to higher neural predictivity.

## 1 Introduction

The fields of machine learning and neuroscience have a long and deeply intertwined history [1]. In the quest for developing intelligent systems capable of learning and thinking by themselves, researchers have repeatedly looked for inspirations in the biological brain. The first generation of Artificial Neural Networks (ANNs) developed in the 1950s utilized perceptrons, which are abstract mathematical models of biological neurons [2]. In the subsequent generations of ANNs, engineering efforts to successfully train these networks eventually led to the design of artificial neuron models that differ from their biological counterparts. Simultaneously, researchers continued to seek and implement biological inspirations for improving ANNs, including their structure and function. For instance, multi-layer convolutional neural networks developed in the 1990s [3],[4] were heavily inspired by the functioning of simple and complex cells in the human visual cortex [5]. More recently, the development of attention networks [6] was motivated by the observation that human brains “attend to” certain parts of inputs when processing large amounts of information.

While the desire to emulate more advanced functions of biological brains serves as one driver of brain-inspiration in the field of ANNs, a second, equally important motivation arises from the need for efficiency. While ANNs have matched or surpassed human performance in many machine learning tasks, including image recognition, machine translation and speech recognition, the computational cost required to do so is quite high and increasing rapidly. Amidst the justified excitement about the success of artificial intelligence in man vs. machine contests such as IBM’s Watson [7] and Google’s AlphaGo [8], the gap in energy efficiency between artificial and natural intelligence continues to grow. Improved energy efficiency is crucial in the face of exploding computational requirements for training state-of-the-art ANNs on the one hand [9], and the need to deploy them in highly energy-constrained energy devices on the other hand [10]. Recent efforts also suggest that biologically inspired mechanisms also have the potential to improve the robustness of ANNs to adversarial attacks [11][12].

Several efforts have explored the use of biologically-inspired concepts for improving the energy efficiency and robustness of ANNs, or allowing them to learn from less data. Among these efforts, one group attempts to increase *representational similarity* at the individual neuronal and synaptic level. For instance, spiking neural networks comprise of neurons mimicking the firing behavior of biological neurons while employing different neural coding schemes [13]. A second group of efforts explore biologically inspired learning rules like Spike-Timing-Dependent Plasticity (STDP) [14]. Finally, other efforts attempt to create ANNs with topologies that are derived from neuro-anatomy [15]. In summary, prior efforts have taken various approaches in the attempt to identify various desirable features of biological brains and embody them in ANNs.

In this work, we focus on quantifying the information similarity between ANNs and biological networks by comparing their internal responses to a given input stimulus [16][17]. This approach was pioneered by Brain-score [16], which quantifies information similarity through a combination of a behavioral sub-score and a neural predictivity sub-score. We specifically focus on *neural predictivity*, which quantifies the ability to predict the responses of a biological brain given the information from an ANN (such as its internal activations), when both are presented with the same stimulus. Brain-Score utilizes the Pearson correlation coefficient to capture the correlation between ANN activations and neural recordings from the macaque visual cortex [16]. The use of Pearson’s correlation coefficient implicitly assumes a linear relationship between the ANN activations and neural responses.

In this work, we advocate the use of an explicit, non-linear mapping function to predict neural responses from ANN activations. The rationale behind this approach is that ANN activations are themselves a product of non-linear transformations. In addition, there does not exist a one-to-one correspondence between ANN and brain layers, decreasing the likelihood that the relationship between ANN activations and neural recordings can be modeled by a linear function. A second key idea that we propose is the use of a neural network to approximate the mapping function itself.

Embodying the approach outlined above, we propose a new method for neural response prediction in order to quantify the informational similarity between an ANN and a set of brain recordings. The method utilizes a neural network, called the neural response predictor (NRP) network, to model the non-linear relationship between ANN activations and brain recordings. Input stimuli (in our case, images) are fed to the ANN and the activations of its layers are extracted. These activations, along with the corresponding neural recordings (captured after presentation of the same stimuli to a primate) [16], are then used to train the NRP network. The prediction error of the NRP network, termed NRP-error, is a quantitative measure of the ANN’s neural predictivity.

We implement the proposed method within the TensorFlow [18] machine learning framework and apply it to calculate the NRP-errors of 8 state-of-the-art image classification ANNs. We utilize neural recordings from the IT (168 neurons) and V4 (88 neurons) regions of primate brains for 3200 images [16] to evaluate the proposed method. Our results demonstrate considerable improvement in neural predictivity over linear models which are used in previous approaches [16]. Our results further indicate that recent advances in image classification ANNs (from AlexNet to Xception) are not accompanied by an improvement in neural predictivity. Finally, we also evaluate the impact of commonly used network optimizations such as pruning on neural predictivity.

## 2 Materials and Methods

In this section, we first describe the general concept of quantifying brain similarity through neural predictivity. We next present the proposed method to quantify neural predictivity and finally discuss the experimental setup and methodology used to evaluate our proposal.

### 2.1 Quantifying Brain Similarity Through Neural Predictivity

Neural predictivity refers to the ability to predict biological neural behavior using the information inside an ANN. As illustrated in Equation 1, one way to quantify neural predictivity is to explicitly formulate a function *f()* that maps ANN activations into predicted neural recordings. In this equation, *Act*_*i*_ refers to the activations of layer *i* in the ANN and *NR*_*pred*_ refers to the predicted neural responses.

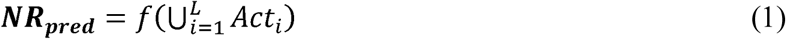

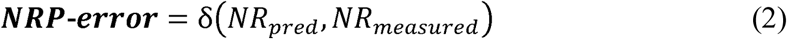

The inputs to the function *f()* are the collection of activations from all or a subset of the layers of the ANN. Next, the predicted neural responses are compared to the actual neural recordings using a distance metric δ such as mean absolute error, to quantify neural response prediction error (*NRP-error*), as illustrated in Equation 2. The NRP-error may be calculated separately for different brain sub-regions (e.g., V1, V4 and IT of the visual cortex) and then averaged to compute the overall error. While there exists a wide range of possibilities for function *f()*, based on the fact that neural networks are universal function approximators, we propose to use a neural network to map from ANN activations to predicted neural recordings.

### 2.2 Neural Response Prediction Method

Our work proposes a new method for quantifying the neural predictivity of an ANN that is based on the overall approach proposed in Section 2.1. The first key idea we propose is to explicitly map ANN activations into predicted neural recordings. A non-linear function is used for this mapping in order to overcome the limitations of previous work [16]. A second key idea is to use a neural network to approximate this non-linear mapping from ANN activations to predicted neural responses.

Figures 1(a) and 1(b) presents the proposed method to quantify the neural predictivity for a given ANN, and given a set of neural recordings. The method consists of the following steps:

- *Transform the ANN by adding the NRP network:* The NRP network is an auxiliary structure that is added to the ANN in order to generate neural response predictions. The structure of the NRP network is detailed in Figure 1(b). First, activations (layer outputs) from selected layers of the ANN are passed through a layer of neurons that we call NRP-L1. The layer NRP-L1 has locally dense connectivity, i.e., the activations from each layer of the ANN are processed separately. This decision was made in order to keep the number of parameters in NRP-L1 and the overall NRP network small. We then concatenate the outputs of NRP-L1 and pass them through one or more dense layers (NRP-L2, …). The final layer in the NRP network (NRP-out) produces the predicted neural recordings. Therefore, the number of outputs of the NRP-out layer is set to be equal to the number of neural recording sites for which data is available. Overall, the NRP network forms a regression network that maps ANN activations into predicted neural responses.
- *Train the NRP network:* The composite network (the original ANN with added NRP layers) is trained while locking down the original ANN’s weights. The training data for this composite network consists of stimuli (images) along with corresponding neural recordings from the visual cortex when the primate was presented with these stimuli. The loss function for this training is the mean squared error between the actual and predicted neural recordings. Standard gradient-based optimizers are used for this step (in our experiments, the Adam optimizer [22] was found to give the best results). A held-out set of data is used to validate the NRP network.
- *Network architecture search for the NRP network:* A key challenge faced by the proposed method arises from the limited number of neural recordings, which translates to limited training data for the NRP network. Although it is reasonable to expect this limitation to be gradually relaxed as additional experiments are performed, it is nevertheless one that must be considered in our effort. Thus, it becomes extremely important to determine an optimized configuration for the NRP network so that it has sufficient modeling capacity to predict the neural recordings, but can also be trained with the limited training data available. We address this challenge by performing a network architecture search [19] on the NRP network. Specifically, we performed a grid search on the following hyperparameters for the NRP network: (i) ANN layers used as input to the NRP network, (ii) sizes of the NRP network layers (except NRP-out, whose outputs must match the number of neural recording sites), and (iii) learning rate.

**Figure 1.**
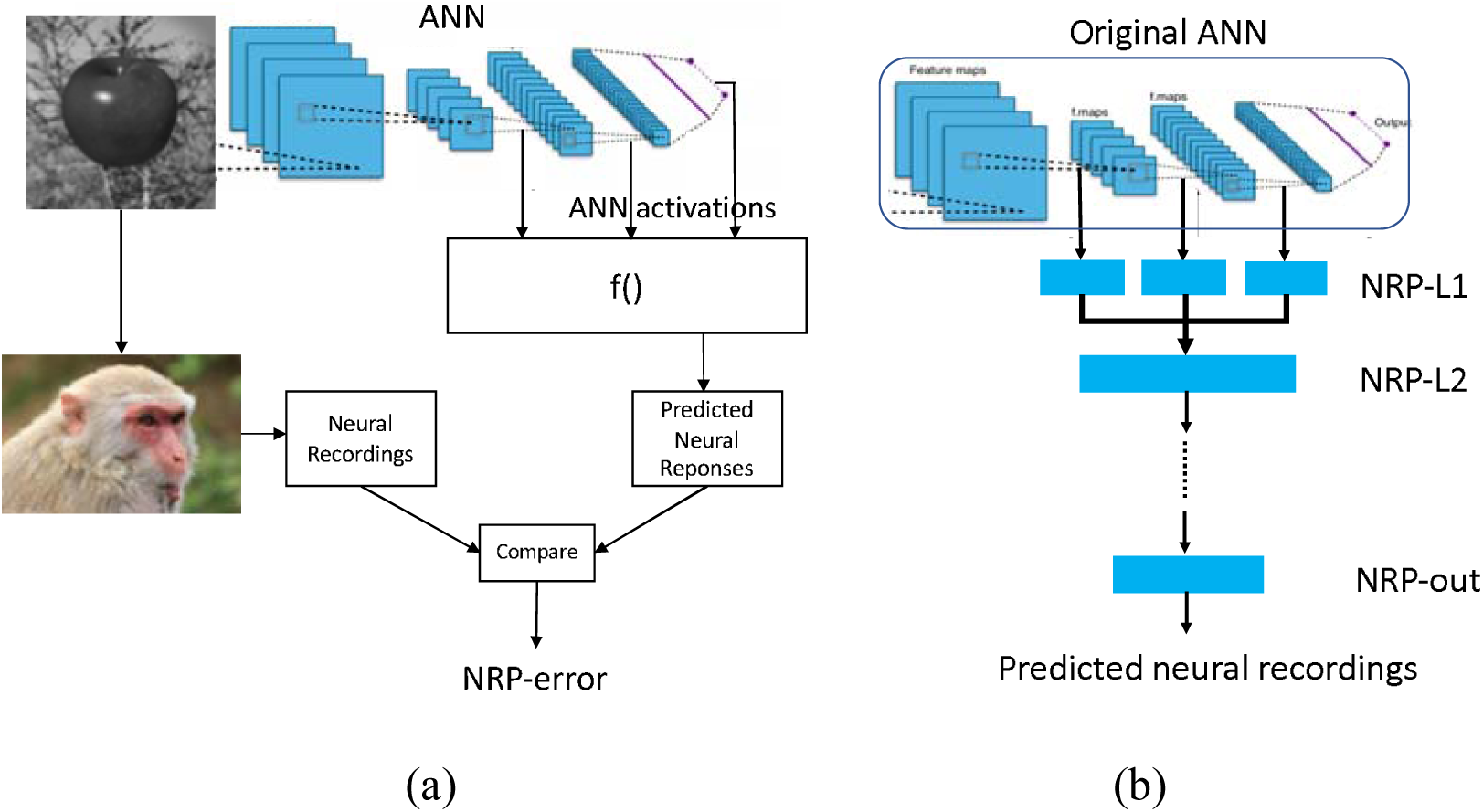
(a) Overview of the proposed method for quantifying neural predictivity, and (b) augmenting an ANN with the Neural Response Prediction network.

### 2.3 Experimental Setup

The proposed method to compute the neural predictivity of an ANN was implemented using the Tensorflow [18] machine learning framework. NRP-errors were calculated for 8 popular image recognition ANNs that have been proposed in recent years for the ImageNet Large Scale Visual Recognition Challenge (ILSVRC) [20]. The characteristics of these networks are described in Table 1.

**Table 1.** Accuracies and NRP-errors of Image Recognition ANNs

| Network | Parameters | Top-5 accuracy | Top-1 accuracy | NRP-error |
| --- | --- | --- | --- | --- |
| MobileNet | 4,253,864 | 0.895 | 0.704 | 0.237 |
| MobileNetV2 | 3,538,984 | 0.901 | 0.713 | 0.249 |
| NASNetMobile | 5,326,716 | 0.919 | 0.744 | 0.247 |
| ResNet50 | 25,636,712 | 0.921 | 0.749 | 0.290 |
| Xception | 22,910,480 | 0.945 | 0.79 | 0.257 |
| DenseNet121 | 8,062,504 | 0.923 | 0.75 | 0.249 |
| DenseNet169 | 14,307,880 | 0.932 | 0.762 | 0.249 |
| AlexNet | 60,954,656 | 0.803 | 0.57 | 0.226 |

The dataset used to train the NRP network and compute NRP-error consists of recordings from 168 neurons in the IT sub-region and 88 neurons in the V4 sub-region of the primate visual cortex [16]. These responses were measured when visual stimuli (3200 images) were presented to the primates (rhesus macaques) for 100 ms each immediately before these measurements were made [16]. Specifically, these neural recordings consist of the average neuronal firing rate for each neuron between 70 ms and 170 ms after the image was presented. Neuronal firing rates were normalized to the firing rates resulting from a blank gray stimulus. Note that the proposed method is generic and can be applied to recordings from additional sites or brain regions as such recordings become available.

NRP-errors were calculated separately for the V4 and IT regions of the visual cortex. In addition to the non-linear model used to generate predicted neural recordings for the calculation of NRP-error, we also implemented a linear regression model to predict neural recordings as a representative of previous efforts.

## 3 Results

In this section, we discuss the results of implementing the proposed method to quantify neural predictivity of ANNs.

Table 1 presents the NRP-errors of 8 different ImageNet classification ANNs. These NRP-errors were computed as the averages of the errors on the V4 and IT regions. As can be seen from the table, some of the more recent ANNs such as ResNet50 (NRP-error of 0.290) are associated with NRP-errors that are higher than older networks such as AlexNet (NRP-errors of 0.226). In fact, AlexNet achieved the lowest NRP-error, while having the lowest Top-1 accuracy, among all networks evaluated. In other words, improvements in application performance (Top-1 accuracy) have not been accompanied by increases in neural predictivity. Another observation is that deeper networks do not necessarily lead to higher neural predictivity. For example, comparing DesneNet121 and DenseNet169, we can see that the additional layers improve Top-1 accuracy but not the neural predictivity. This overall trend, illustrated in Figure 2(a), is consistent with observations from recent efforts on quantifying brain similarity [20]. This is perhaps because, deeper ANNs have enabled improvements in accuracy, but have done so by adopting internal representations that are beyond and less like those used in biological systems.

**Figure 2.**
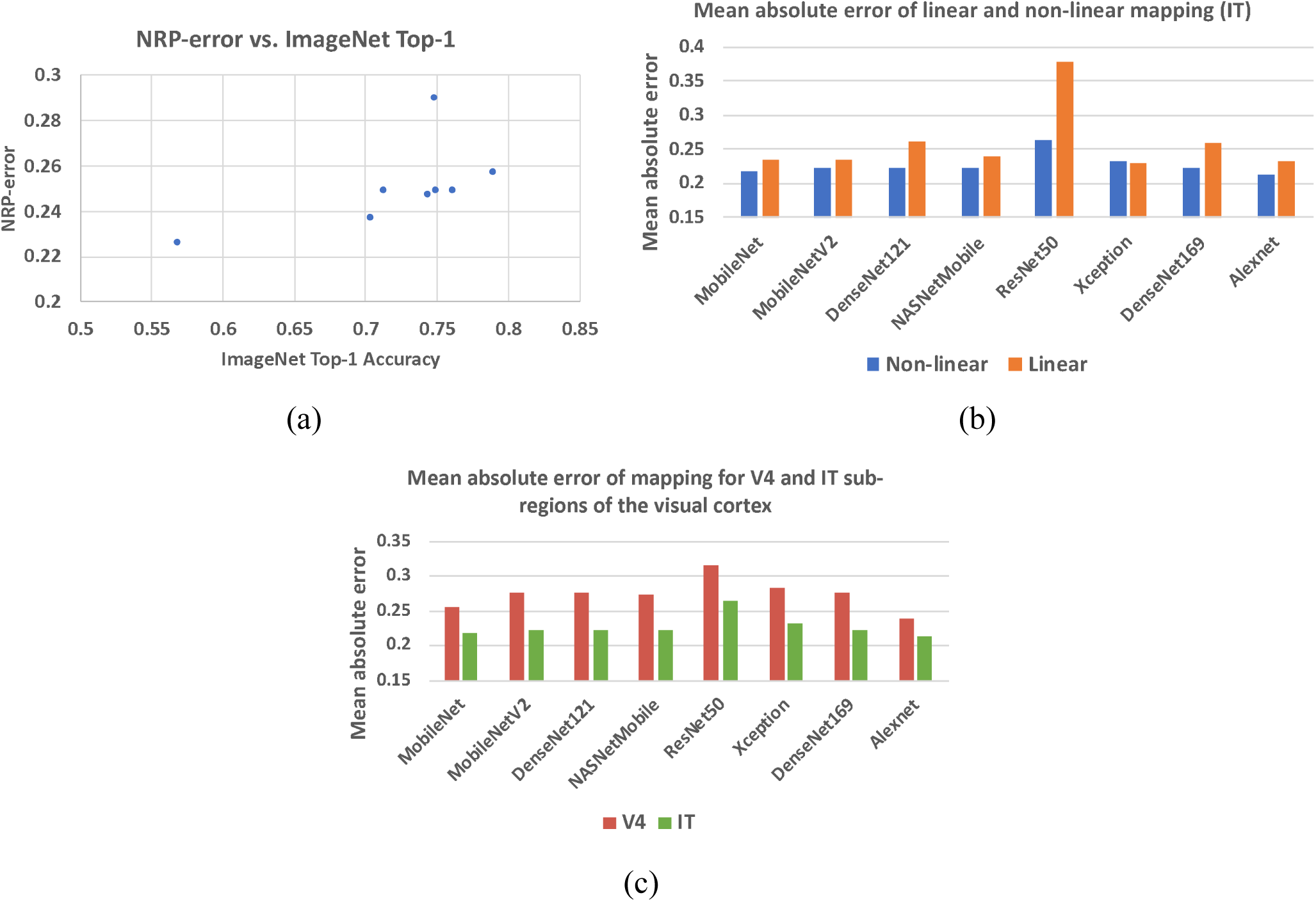
(a) NRP-error vs. Top-1 accuracy, (b) neural response prediction error of proposed (non-linear) and baseline (linear) methods, and (c) prediction error for V4 and IT regions.

### 3.1 Necessity of Non-Linear Mapping function

A key feature of our work is the use of a non-linear mapping function (approximated by a neural network) to map ANN activations to predicted neural recordings in the calculation of NRP-error. This is in contrast to prior efforts, which use the Pearson correlation coefficient, effectively assuming a linear relationship between ANN activations and neural responses. In order to demonstrate the necessity of a non-linear mapping function, we also implemented a linear regression model to predict neural recordings from ANN layer activations. Figure 2(b) compares the mean absolute error obtained from the proposed method as well as the linear regression model for the IT region. As can be seen from Figure 2(b), our results show that a non-linear mapping function from ANN activations to predicted neural recordings significantly decreases the error of neural prediction and can hence be considered a superior predictor of an ANN’s neural predictivity. The results for V4 also lead to the same conclusion. For example, in the case of Resnet-50, the mean absolute error of the linear and non-linear models are 0.379 and 0.265, respectively. This is explained by the facts that ANN layers are non-linear transformations and there is no layer-to-layer correspondence between most ANN and brain layers, making a non-linear function more suitable to model the mapping between ANN activations and neural recordings.

### 3.2 NRP-errors for V4 and IT sub-regions

In order to compare the neural predictivities for the V4 and IT sub-regions of the visual cortex, NRP-errors were computed separately for both sub-regions. From the results, we observe that ANN activations predict IT neural recordings with a higher accuracy than V4 neural recordings (Figure 2(c)). This suggests that ANNs use representations that have a higher level of similarity with later visual cortex sub-regions (such as IT).

### 3.3 Relationship Between Neural Predictivity and Layer Sizes of NRP Network

To overcome the small amount of training data (neural recordings) available for the NRP network, a suitable configuration must be determined so that it has sufficient modeling capacity to predict the neural recordings but can also be trained with the limited training data available. In order to address this, we performed a network architecture search on the NRP network by varying the sizes and number of intermediate layers to find a suitable configuration. Representative results obtained for the MobileNet ANN are presented in Figure 3(a). We found that using two intermediate layers in the NRP network before the output layer is sufficient to model the mapping from ANN activations to predicted neural recordings. We also found that there is a sweet-spot of layer sizes for the NRP network that minimizes the average mean absolute error of mapping across all networks.

**Figure 3.**
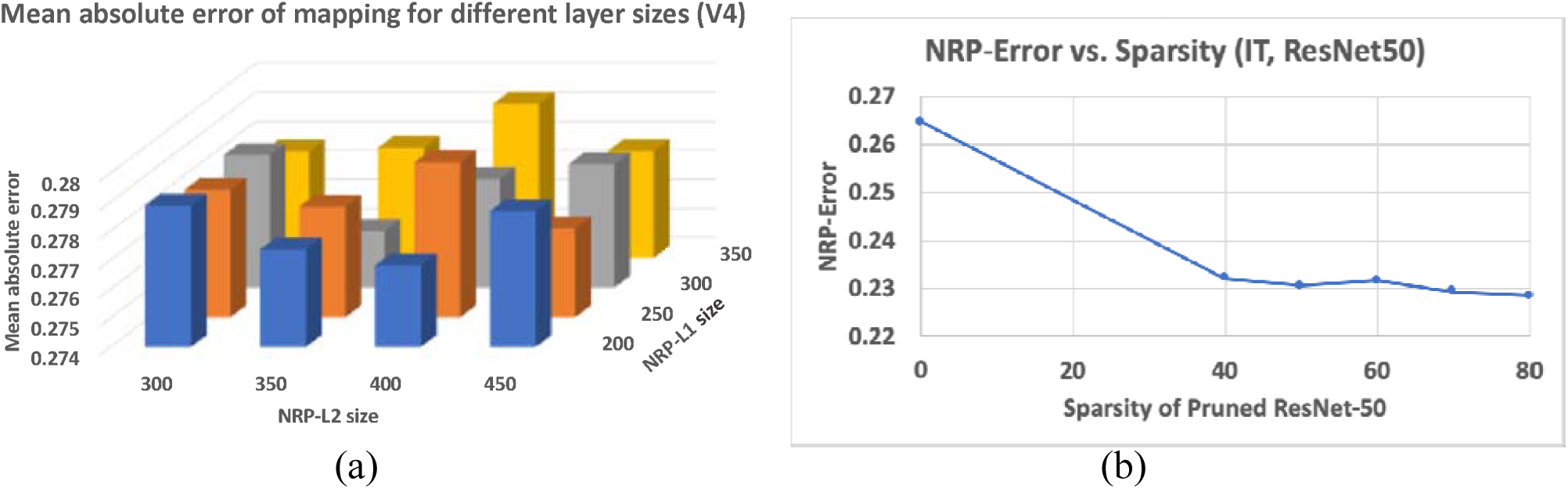
(a) Impact of NRP network layer sizes on prediction error, (b) impact of network pruning on NRP-error.

### 3.4 Impact of Network Pruning on Neural Predictivity

Finally, we investigate the impact of a popular ANN optimization technique, namely network pruning, on neural predictivity. We consider the ResNet50 ANN and applied state-of-the-art pruning algorithms [21] to derive pruned models with varying levels of sparsity. We then applied the proposed method to the pruned models to compute the corresponding NRP-errors, and the results are presented in Figure 3(b). The results suggest that pruning has a clear decrease in NRP-error, indicating a positive effect on neural predictivity. We believe this is due to the fact that pruning removes “extraneous” information from the ANN, making it easier to map its activations to the neural recordings.

## 4 Discussion

Despite the rapid advances made in the field of deep learning over the past decade, biological brains still have much to teach us in the quest to build more energy-efficient and robust artificial intelligence. A key step towards drawing inspiration from biological brains is to quantify the similarity between them and their artificial counterparts. Our work takes the approach of quantifying similarity through neural predictivity, or the ability to predict neural responses from a biological brain given the internal information of ANNs. Since this is the goal of our work, we discuss closely related efforts and place our own effort in their context. We also discuss possible future directions, both in terms of improving our work and its applications.

### 4.1 Related Work

A recent effort that quantifies neural predictivity is Brain-Score [16]. Brain-Score specifically focuses on evaluating ANNs that perform core object recognition tasks, and provides a quantitative framework to compare image classification ANNs with measurements from the visual cortex of primates (firing rates for specific neurons when the primate is presented with the stimulus). It consists of a behavior sub-score and neural predictivity sub-scores for various regions of the visual cortex (V1, V2, V4, IT). The behavior sub-score quantifies how similar the ANN’s predictions are to those made by the primate when both are presented with the same stimulus. The neural predictivity sub-scores capture how well the ANN’s activations correlate to the neural recordings from each region of the visual cortex. These sub-scores are computed as the Pearson correlation coefficient between ANN layer outputs and neural firing rates for that region.

Through the use of the Pearson correlation coefficient, Brain-Score implicitly assumes a linear relationship between ANN activations and neural firing rates. However, since ANN layers are non-linear transformations, there is no evidence to support this assumption. Moreover, there is no layer-to-layer correspondence between most ANN and brain layers, making the likelihood of a linear relationship even less likely.

Our work extends the state-of-the-art through two key ideas. First, it advocates the use of an explicit (non-linear) mapping function to predict neural responses from ANN activations in order to quantify neural predictivity. A second key idea is the use of a neural network (known to be a universal function approximator) to approximate the mapping function itself. Our experiments clearly support the merit of these proposals by demonstrating an improved ability to predict neural responses.

### 4.2 Future Work

One possible direction to build upon our effort would be to collect and incorporate additional neural recordings into the dataset used. A dataset with additional recording locations and more input images would allow us to train larger (and potentially more accurate) NRP networks without the risk of over-fitting. Since internal representations are greatly influenced by training, it would also be interesting to study whether networks trained with bio-plausible learning rules (e.g., STDP) yield higher neural predictivity than ANNs trained with gradient-descent. Finally, building upon a recent result that using brain-like representations in the early layers of an ANN can lead to higher robustness, it would be interesting to study whether there is a relationship between an ANN’s neural predictivity and its robustness to noise and adversarial perturbations.

## 5 Acknowledgments

This work was supported in part by C-BRIC, one of six centers in JUMP, a Semiconductor Research Corporation (SRC) program sponsored by DARPA.

The authors gratefully acknowledge Martin Schrimpf and James DiCarlo from the Department of Brain and Cognitive Sciences at the Massachusetts Institute of Technology for providing them with the neural recordings used in this work, and for their valuable suggestions.

## Notes

### Competing Interest Statement

The authors have declared no competing interest.

